# Whose energy cost would birds like to save? a revisit of the migratory formation flight

**DOI:** 10.1101/2023.03.17.533072

**Authors:** Mingming Shi, Ignace Ransquin, Philippe Chatelain, Julien M. Hendrickx

## Abstract

Line formation of migrating birds is well-accepted to be caused by birds exploiting wake benefits to save energy expenditure. A flying bird generates wingtip trailing vortices that stir the surrounding air upward and downward, and the following bird can get a free supportive lift when positioned at the upward airflow region. However, little to no attention has been paid to clarifying birds’ interests in energy saving, namely, do birds intend to reduce their individual energy consumption or the total energy of the flock? Here, by explicitly considering birds’ interests, we employ a modified fixed-wing wake model that includes the wake dissipation to numerically reexamine the energy saving mechanism in line formation. Surprisingly, our computations show that line formation cannot be explained simply by energy optimization. This remains true whether birds are selfish or cooperative. However, line formations may be explained by strategies optimizing energy cost and either avoiding collision or maintaining vision comfort. We also find that the total wake benefit of the formation attained by selfish birds does not differ much from that got by cooperative birds, the maximum that birds can attain. This implies that selfish birds are still able to fly in formation with very high efficiency of energy saving. In addition, we explore the hypothesis that birds are empathetic and would like to optimize their own energy cost and the neighbors’. Our analysis shows that if birds are more empathetic, the resulting line formation shape deviates more from a straight line, and the flock enjoys higher total wake benefit.

**Author summary:** Migratory birds can achieve remarkable performance and efficiency in energy exploitation during annual round-trip migration flight. Theoretical and experimental results have shown that this might be achieved because birds fly together in formation with specific shapes, e.g. the noticeable V formation, to utilize the aerodynamic benefits generated by their flock mates. However, it is still unclear whether energy-guided behavior indeed can lead to these formations. We show that the special formation adopted by migratory birds cannot be explained purely by the energy exploitation mechanism, and that birds’ vision performance and collision avoidance very likely also play important roles in the formation emergence. Our results imply that birds fly together in formation because of energy saving, but the specific shape of the formation depends on non-aerodynamic reasons. The research provides further understandings of the emergence of migratory formation and the energy saving mechanism of animal groups. It may also indicate that wing flapping, currently not considered, has an important effect on the way birds exploit aerodynamic benefits from others during the formation flight.

---

Highly organized motion behaviors are commonly observed among animals in nature. Researchers have been long fascinated by the emergence of these collective motions, which could be caused by reduction of energy expenditure, enhancement of motion performance and social comfort [10, 20, 21]. As for bird migrations, the *line formation* [15], including the echelon, J and V shapes, has been long observed [14, 15], although not for every migratory species. Systematic explanation of these formation shapes attracts attention of scientists from biology, aerodynamics, mechanics and computer science. It is widely accepted that line formation can reduce birds energy cost [12, 15–18, 28–37], which is critical for birds survival in long-distance and sustained migration journeys [18, 46]. Theoretically, the fixed-wing aerodynamic wake model predicts that a flying bird generates a pair of symmetric trailing vortices downstream, slightly inside of the wingtips. The vortices induce upward airflow outboard of the wingtips. Hence the following bird gains free lift and reduces its drag when riding on the upwash of the front bird [31, 34]. We note that this wake model does not take wing flapping [17] into account but remains valid as a first approximation, and accurately captures gliding birds and flights with low flapping rate, typically occurring for large size migration birds, e.g., swan and geese [21]. The energy cost reduction predicted by the aerodynamics theory was validated experimentally for both aircraft formation [2, 3] and birds formation [33], though indirectly for the latter: the measured heart rate and wing beat rate of the great pelicans flying as followers in the formation are smaller than those in solo flight, corresponding to energy savings of 11-14%. Other tests [35, 36, 38, 46] also observed that the lateral distance of a bird to its front neighbor in the formation also fits with the theoretical result from the fixed-wing aerodynamic wake model, and therefore indirectly proves the energy saving of line formations.

Apart from energy saving, it is long debated that formation flight of migration birds may also occur to enhance the visual contact [14–16, 28–31, 37]. It is mentioned in [39, 40] that the density of single and double cones on a bird’s retina, which is correlated with the visual performance, is highest around the vision axis (fovea). Heppner speculated from experiment [37] that Canada geese have nearly immobilized eyes. Hence to achieve the highest resolution and mitigate vision discomfort, it is best that the front neighboring bird locates around the back bird’s fovea. The enhancement of visual contact could also enable the young bird to learn the migratory path and behavior of senior birds, benefiting the migratory group. The visual factor could work as a complement to the aerodynamic wake benefit in leading the formation [22, 28, 31]. However, it may be less important in determining the formation shape, as the angles of the V formation predicted by Heppner [37] are out of the range of the observations [27] and the correlation between the lateral and longitudinal distances of neighboring birds is weak [16, 28, 36]. On the other hand, one can expect that, avoiding colliding with other birds when birds are close must be a critical factor that affects birds transient trajectories or even the formation configuration [11, 21, 24]. Indeed, two birds cannot overlap physically. Moreover, the stress of collision risk when birds are very close could raise their metabolic level [21], increasing birds energy consumption, or in other words reducing their wake benefits.

Although many works were dedicated to mechanisms leading to the emergence of bird line formations, very few explicitly paid attention to birds interests in formation: whose energy expenditure birds would like to save. It is natural to assume that each bird is selfish and only saves its own energy consumption. However, selfishness can harm the whole group, with the prisoner dilemma as a typical example. Hence, to maximize the whole group survival, migratory birds may have evolved after several million years to behave somehow collaboratively in order to save the energy expenditure of the entire flock. Some works have unconsciously adopted the selfish bird assumption [16, 28, 34–36, 45]. However, the longitudinal and/or lateral distances to the front neighbor of the bird in the formation are often decided by assuming implicitly that each bird only exploits the upwash from the front neighbor. This is unrealistic since every bird is immersed in the wake field induced by the vortices of all other birds.

Though the wake from distant birds might be neglected, it is argued that a bird can still get wake benefit from the back neighbor [22, 30, 31]. As for cooperative birds, even though many papers took the energy saving of the group as the flight performance index, very few discussed the optimal formation shape. This might be because the wake model used in these works ignores the slow decay of the vortex strength [19, 42–44], and predicts thus that the total energy saving is unchanged when birds slide in the flight path, so that a wide range of formations are equivalent in terms of energy savings.

However, the wake dissipation may overturn this conclusion. Although [45] has noticed the wake dissipation, the Gaussian decay model used there for the vortex dissipation makes the vortex strength decays too fast with respect to experimental results [49], which may have an important impact on their results. Moreover, [45] omitted the bound vortex [22, 31, 32] that may also be important in the near field of the bird, and assumed the same optimal relative position of each bird to its front bird. Since bird interests in energy savings straightforwardly influence their behavior, the neglect of bird interests in the existing literature renders the conclusion that birds adopt the line formation because of energy saving questionable.

In view of these, the actual reasons behind the emergence of line formations remain unexplained, and several important questions arise naturally: does the line formation emerge due to migrating birds saving energy expenditure following a certain behavior pattern? If not, should other factors, like visual contact and/or collision avoidance also play roles in determining the formation configuration? Could birds only behave selfishly in formation flight, or is cooperation necessary for birds in forming the line formation. To answer these questions, we examine how the combination of aerodynamic wake benefit, visual contact or collision avoidance with bird interests in energy saving, determines the emergence of the line formation. This is accomplished by testing if there exist stable bird position configurations that correspond to line formations and maximize birds wake benefits under different behavior assumptions and/or non-aerodynamic factors. Our wake model is based on the most basic and widely used horseshoe model [22, 28, 31–35, 45] and also characterizes the slower vortex dissipation in the close formation, closer to the real wake data [44, 49]. Moreover, our wake-induced model is accurate in the sense that each bird is affected by the wake induced by all other birds. In this situation, it is unclear whether the line formation assumed to have the same relative position of neighboring birds [16, 22, 28, 31, 36] could appear.

Surprisingly, our results show that line formations cannot emerge purely based on the energy saving mechanism, for both selfish and cooperative birds. For selfish birds, there exists no stable formation configuration that birds can maintain, let alone the line formation. This is mainly due to the fact that a bird can gain marginally more wake benefit from following birds than that from leading birds if it drifts down stream. For cooperative birds, the stable formation configuration exists but results in collisions. By contrast, the results show that line formation could be explained by birds maximizing their own energy saving and maintaining vision comfort. While for cooperative birds, the results show that the effort to avoid collision forces birds to be separated when they reduce group energy expenditure, also leading to typically observed line formations. Our results also show that the total wake benefit of selfish birds in the formed line formation only reduces slightly (by around 5%) compared with that of cooperative birds. Since the aerodynamic wake benefit is quite complex, we speculate that birds mostly behave selfishly in forming the line formation, in consideration of the trade-off between the complexity of information processing and energy efficiency. Finally, taking into account both vision performance and collision avoidance, we focus on an intermediate situation between selfish birds and cooperative birds, where birds show empathy by also taking into account the wake benefit of their neighbors. We show that if birds are more empathetic, the formation shape is close to that of cooperative birds with more variability, though the flock achieves higher energy efficiency.

## Results

### Model

We consider a flock of *n* + 1 birds in a steady-state horizontal formation flight on the same plane: the flight direction, height and velocity of the formation remain constant. We focus on the left echelon formation, where each bird is positioned behind and to the left of its front neighbor, since echelon is the most common shape [14, 21]. We assume that birds have the same size and weight and nearly fly in an echelon shape. The birds are ordered from front to back, with the leading bird labeled 0, and the *i*th closest bird to the leader labeled *i*. Two birds are neighbors if the difference of their order is one. Since only relative positions matter in the formation shape, we set the origin of our frame of reference at the center of the leader. The *x* axis of the frame is chosen towards the flying direction, and the *y* axis points in the right-hand direction, perpendicular to the *x* axis. The position of bird *i* in the frame is denoted as *p*_*i*_ = [*x*_*i*_ *y*_*i*_]^⊤^, then *p*_0_ = [0 0]^⊤^. Since we assume that the flock is close to an echelon shape, *x*_*i*_ ≤ *x*_*i*−1_ and *y*_*i*_ ≤ *y*_*i*−1_ for each *i* = 1, 2, …, *n*.

As aforementioned, migration birds flying in a line formation may lower their energy consumption by exploiting the wake phenomenon: the flight of a front bird stirs the surrounding air upward and downward. The birds that position themselves properly relative to their predecessor can gain beneficial free lift from the upward airflow, leading to drag reduction. We treat each bird as a fixed wing and characterize the wake benefit by the supportive airflow velocity. The latter is extended by a modified horse-shoe vortex model, including the phenomenon of slow wake circulation strength decay [44]. In detail, the wake of a constant-velocity wing consists of a finite bound vortex around the wing and two semi-infinite vortices that start at the wingtips and extend downstream. The vertical airflow induced by the vortices points downward right behind the bird and upward in the region outboard from the wingtips. Note that there also exists upward airflow at the front of the wing. For a bird *i*, we take the averaged vertical airflow velocity over its wing induced by the wake of another bird *j* as the wake benefit *f* (*p*_*ij*_), with *p*_*ij*_ = *p*_*i*_ − *p*_*j*_ the relative position of bird *i* to *j* [22, 31, 45] (See Method for details). Figure 1 shows the function *f* (·). The wake benefit *f*_*i*_(*p*) of a bird *i* from all other birds *j* in the flock, with *p* the vector of all birds’ positions, is assumed to be the sum of *f* (*p*_*ij*_). Note that we neglect the sidewash and the rolling moment on the bird induced by the non-even distribution of the vertical air velocity from the wake of another bird [19], which is realistic according to [22].

**Fig 1.**
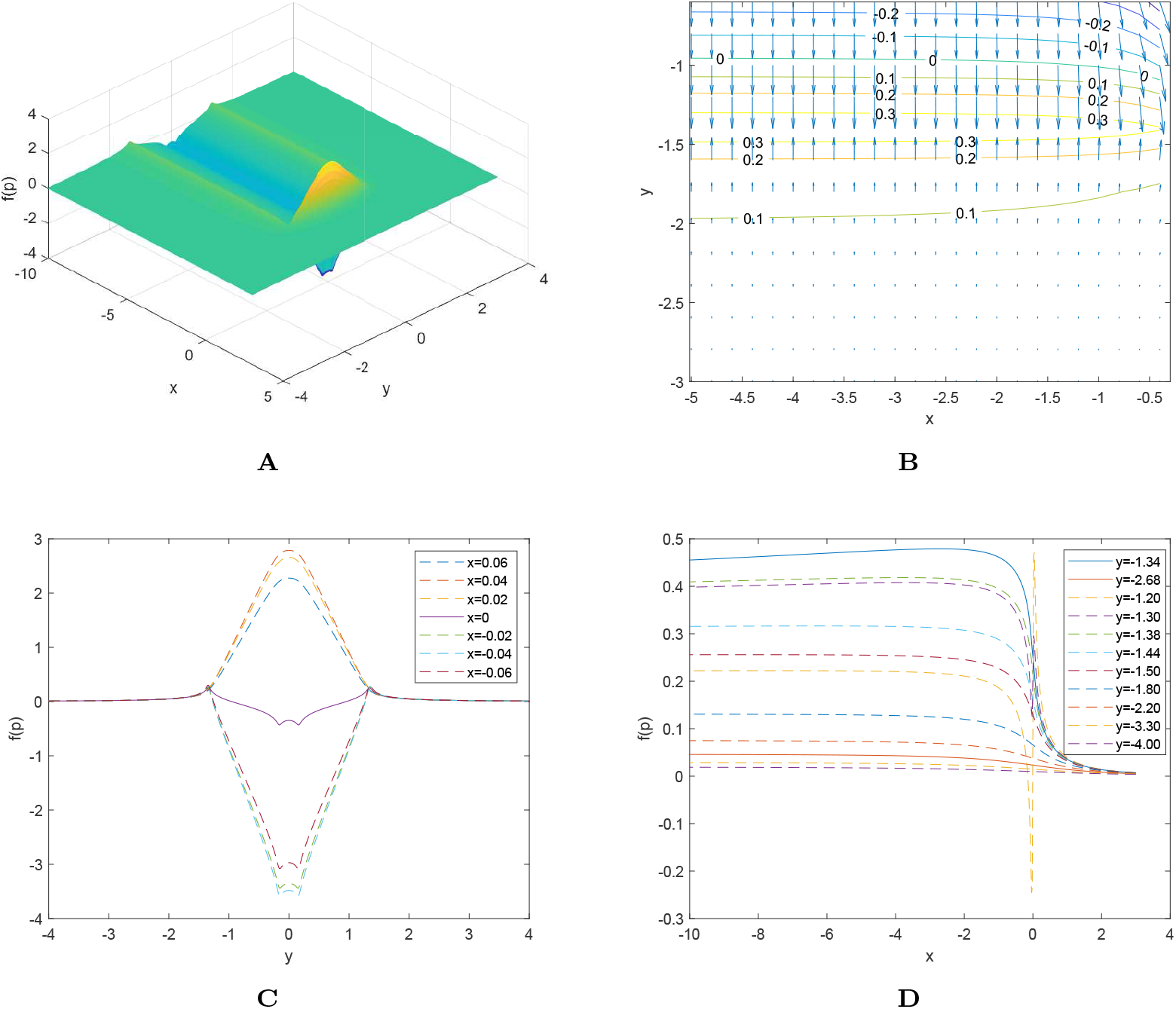
The aerodynamic wake benefit by the horse-shoe wake model. Two birds with same size and weight are flying at the *x* direction. The figures show the wake benefit *f* (*p*) induced by the first bird at the origin on the second bird at the position *p* = [*x y*]^⊤^. The parameters for birds and the wake model are set as *b* = 1.5m, *r*_*c*_(0) = 0.02*b, W* = 36.75N, *ρ* = 1.112kg/m^3^, *U* = 18m/s, and *D*_*f*_ = 0.0014m^2^*/*s (Selected for Canada geese [9] and see Method for notations). The wake benefit function *f* (*p*) peaks slightly before the first bird. After the first bird, it peaks around two lines *y* = *y*^∗^ = − 1.340 and *y* = − *y*^∗^ = 1.340. More specifically, in the fourth quadrant ((−∞, 0) × (−∞, 0)), around the line *y* = *y*^∗^ and along the streamwise direction (negative *x* direction), *f* (*p*) increases quickly to maximum, then decays very slowly. A slight deviation of the *y* coordinate from *y*^∗^ leads to a large decrease of wake benefit. When |*y*| ≥ |*y*^∗^|, the larger the lateral distance of birds is, the flatter *f* (*p*) is along the *x* direction, and the smaller the *x* that makes *f* (*p*) peak is. Also if |*y*| ≥ |2*y*^∗^|, the wake benefit of the second bird varies little when its lateral distance to the first bird changes. The maximum of *f* (*p*) in the fourth quadrant is (*x*^∗^, *y*^∗^) = (− 2.601, − 1.340). **A**: The wake benefit *f* (*x, y*). **B**: Contour of *f* (*p*) and gradient field in the vicinity of the leader. The arrow represents the local direction at which the wake benefit function increases. **C**: Slice of wake benefit *f* (*x, y*)-Front view. **D**: Slice of wake benefit *f* (*x, y*)-Left view for negative *y*.

Birds can behave selfishly or cooperatively. When birds are selfish, each follower optimizes its position to maximize the wake benefit *f*_*i*_(*p*). This can be viewed as a game, and we are interested in the Nash equilibrium, corresponding to the stationary formation configuration where every bird’s wake benefit could not increase more if the bird deviates to another position unilaterally. When birds are fully cooperative, the followers are interested in optimizing their positions in a collaborative manner such that the total wake benefit *J*(*p*), the sum of the wake benefit *f*_*i*_(*p*) of all birds (including the leader), are maximized. Birds could also show empathy to the birds close to them; this constitutes an intermediate between purely selfish and fully cooperative. In this situation, each follower may optimize not just its wake benefit, but also the neighbors’. In the later, we will first elaborate on selfish and cooperative birds, then discuss the empathy situation shortly.

We also examine the effect of collision avoidance in the generation of a formation. This is very relevant in our context as energy-guided behavior may in some cases lead to very small bird separations that actually constitute collisions. We model each bird as an ellipse with the semi-major axis *b*_*l*_ and the semi-minor axis *b*_*s*_, which align with the *y* and *x* axis, respectively. See Fig 2**A**. The value of *b*_*l*_ and *b*_*s*_ depend on the wingspan and bill-to-tail distance of the bird. As shown in Figure 2**B**, to avoid the collision with bird *j*, the center of bird *i* must be outside the orange ellipse, whose boundary sets a barrier of bird *j* to other birds. Others should realize the possible future collision, hence a yellow region, slightly enlarged from the orange ellipse, is used to represent the collision alert region of bird *j*. We characterize the barrier and the collision alert region by a potential function *B*_*c*_(*p*_*ij*_) (see Method for details), which satisfies several properties: a) *B*_*c*_ = 0 if *p*_*ij*_ is outside the collision alert region; b) *B*_*c*_ decreases monotonically to −∞ if *p*_*ij*_ approaches the boundary of bird *j*. The barrier function *B*_*c*_ should be built for each pair of birds.

**Fig 2.**
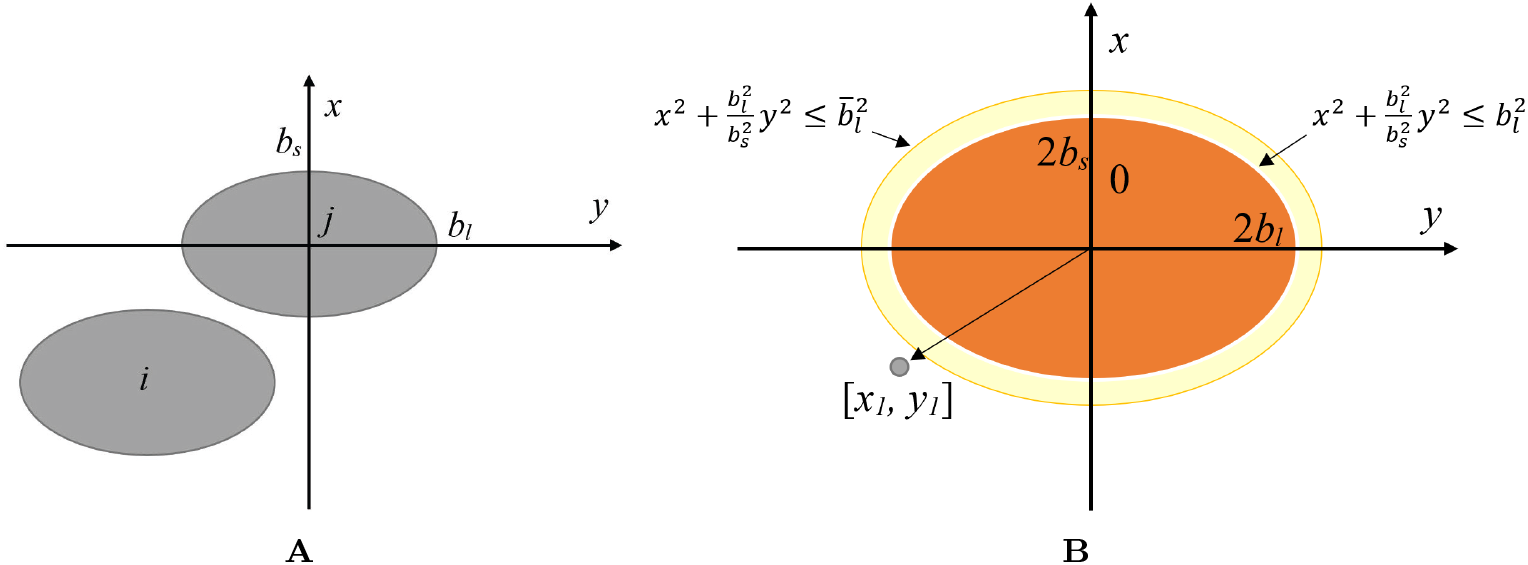
Two birds and the barrier region. **A**: Two birds represented by the ellipses. **B**: The barrier region and collision alert region of bird *j* to another bird *i*. The orange ellipse represents the barrier of bird *j*. If the center of any other bird *i* locates within this region, then *i* collide with *j*. Hence this should be avoided in any case. The yellow region represents the collision alert region. If the center of bird *i* locates within this region, it is aware that a collision with bird *j* can happen if not moving properly.

We also test whether visual communication influences the formation emergence. As argued in [37], the bird may need to align its front neighbor around one of the vision axes (see Figure 3**A**) to avoid eye discomfort. This requirement seems too restrictive, since the cone cells, which determines the vision performance, does not fade abruptly around the fovea area [39, 40]. Hence we assume conceptually that each bird eye has a vision comfort zone (see Fig. 3**A**), which is a sector with spanning angle 2*θ >* 0, centered at the eye and evenly divided by the vision axis. If the front neighbor of a bird locates within the vision comfort zone, the bird feels no eye discomfort, therefore behaves normally. However, if the front birds locates outside of this sector, then the bird feels the vision distortion and tries to eliminate it by repositioning. To characterize the vision comfort, as shown in Figure 3**B**, we assume that the bird eyes locate at the front end of the semi-minor axis of the ellipse that represents the bird body, neglecting the bill-to-eye and the eye-to-eye distance of a bird, since they are often small compared to wingspan and bill-to-tail distance (For Canada goose, the ratios of the eye-to-eye distance to the wingspan and the bill-to-eye distance to the bill-to-tail distance are 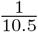 and 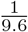, respectively) and only have effects on final results if birds are too close where the formation is inconsistent with observations. We assume that the bird takes the center of another bird as the position of that bird. The vision comfort is quantified by a function *B*_*v*_(*ϕ*_*ij*_), where *ϕ*_*ij*_ is the vision angle of bird *j* in bird *i*’s eyes. The function *B*_*v*_(*ϕ*_*ij*_) is constructed such that if |*ϕ*_*ij*_| ≤ *θ, B*_*c*_(*ϕ*_*ij*_) = 0, implying no vision discomfort, while if |*ϕ*_*ij*_| *> θ* and gets larger, the function *B*_*v*_(*ϕ*_*ij*_) becomes more negative (see Method for details). Although we allow *B*_*v*_(*ϕ*_*ij*_) to approach −∞, this scenario cannot happen in the current setting as explained in Method. We associate to each follower a vision comfort function. During the long-term formation flight, the follower adjusts its position to maximize the vision comfort function. The vision angle can be calculated from the relative position of a bird to its front neighbor, hence *B*_*v*_ is also the function of the relative position of two neighboring birds. In the paper, the parameters for vision angle are selected artificially or partly from Canadian goose [37]. Note that other meticulous models for the vision comfort could also be possible, but the main effects remain the same.

**Fig 3.**
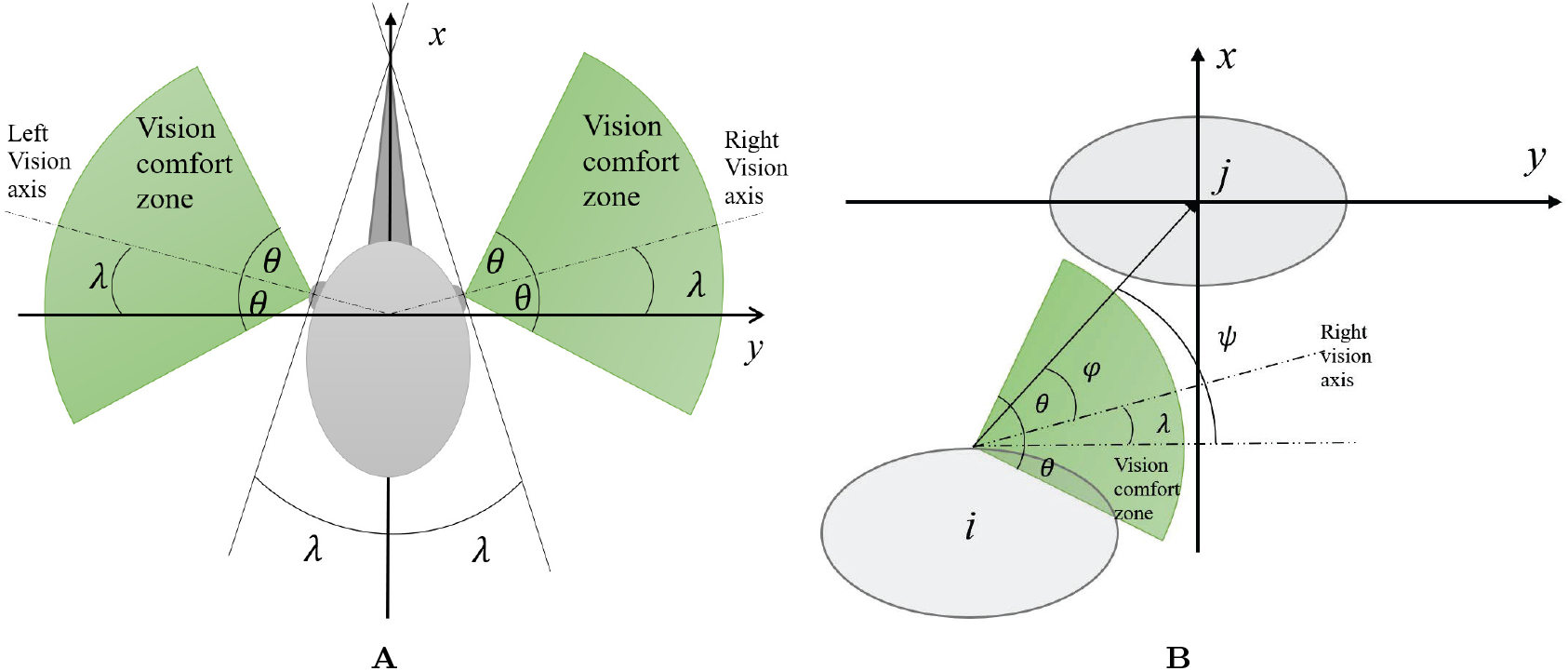
The vision comfort zone of a bird. **A**: Bird head. The variable *λ* is the front binocular vision angle, and 2*θ* represents the vision comfort zone. The left (right) axis is assumed to be perpendicular to the left (right) line that forming the front binocular vision. when an object locates out of the vision comfort zone (green zone), the bird feels the vision discomfort. **B**: Two birds and the right vision comfort zone. If bird *j* locates outside green zone, bird *i* will try to reposition to make bird *j* close to this zone.

We examine whether bird formations emerge from birds optimizing their wake benefits by checking if there exists some stationary point for the game or the optimization problem of maximizing wake benefits. When birds are selfish, the *egoistic equilibrium* designates the Nash equilibrium of the game of birds maximizing *f*_*i*_(*p*). When birds collaborate to maximize *J*(*p*), the solution of the optimization problem is called the *cooperative equilibrium*. An equilibrium can be unstable or marginally stable; birds can thus hardly maintain the corresponding formation shape due to the pervasive disturbances, e.g., wind or perception noise. Hence, we are focusing specifically on the stable equilibria that correspond to line formations, and not interested in others. We ignore how birds adjust their positions to reach the equilibria, but we can at least compute the equilibria (when they exist) numerically. In this paper, we use a gradient-based method to detect the existence of such equilibria. Starting from multiple initial points, we iteratively update the estimate vector *p*^*k*+1^ ∈ ℝ^2*n*^ of the equilibria along the gradient of the local wake benefit and the total wake benefit, respectively, based on the current estimate *p*^*k*^. Let us recall that gradient is popular in optimization problems [41] and also the computation of equilibria of games [1, 4]. It represents what birds would do naturally if they could feel the gradient of the individual or the total wake benefit, which probably happens in the selfish situation since the upwash gradient is proportional to the rolling and pitching moments felt by the bird. Even so, we note that birds do not need to follow the gradient of wake benefits in actual formation flight. Additionally, there may exist multiple equlibria and some are missed in our computation. However, since multiple different initial bird positions are used in our computations, the stability margin of the potential equilibria should be small and such formation would then be easily devastated by noise and disturbances. We also examine whether the stable equilibria exist if birds maximize wake benefit with or without including collision avoidance or/and vision comfort. This is done by adding the gradient of the barrier function *B*_*c*_(*p*) or/and the vision comfort function *B*_*v*_(*p*) to the gradient-based search. Throughout the paper, the parameters related to the wake model are taken from Canada geese [9]. However, this does not imply that our conclusion cannot hold for other birds. In fact, as shown in Method, the scaled optimal positions of birds (the ratio of bird positions over the wingspan) obtained from the pure wake benefit maximization do not change with the variation of bird parameters.

We focus on whether there exists a stable line formation that is energy optimal. There are multiple mechanisms by which birds could achieve such a formation apart from gradient guided action. For instance, optimal behavior could be learned over time, evolution, social pressure etc. [5–7], but we do not consider these here. Also the assumption of fixed leader position seems to violate the selfishness of birds, since as shown later the leader can get more wake benefit if it is closer to the followers. However, assuming that the leader also optimizes its wake benefit by adjusting position would reduce the whole flock speed gradually to zero, an unrealistic situation. On the other hand, it has been recognized that during migration flight, birds often take turns in leading formation [46]. Hence the current leader can also enjoy the wake benefit from other birds in the future. Furthermore, the situation that parenting birds always take the leading position has been observed [8]. In the paper, we do not address the question that how selfish birds switch positions to maximize their long-term wake benefits but rather than focus on the optimal formation at a longer time scale over which no birds switch position. Finally, we stress that no specific kinematics of birds is considered. Hence the bird positions at the obtained equilibria can be interpreted as the limit for more realistic bird kinematics models.

### Insufficiency of energy optimization in formation emergence

Surprisingly, we find that energy optimization alone is insufficient to explain the emergence of realistic line formation, for both selfish and cooperative birds. The simulation shows that no stable egoistic equilibrium for selfish birds exists and the cooperative equilibrium found for collaborative birds is inconsistent with empirical observations. In this subsection, we only present the situation of one leader and two followers, since the results for more followers lead to identical conclusions.

For the case of selfish birds, the non-cooperative game has no equilibrium within the valid zone of the wake model. This is reflected as the non-convergence of the gradient-based algorithm for multiple different initial bird positions: starting from an initial condition, the lateral positions of birds 1 and 2 converge respectively to almost *y*^∗^ and 2*y*^∗^ with *y*^∗^ given in Fig 1, but their longitudinal positions become farther away from the leader as iterations continues, as shown in Fig. 4. In other words, bird 1 and 2 first converge to the lines *y* = *y*^∗^ and *y* = 2*y*^∗^, respectively, then drift backward along these two lines. The absence of a stable equilibrium and the constantly drifting downstream of birds 1 and 2 can be explained qualitatively as follows. On these two lines, the wake benefit of birds 1 and 2 comes mainly from their front neighbor, bird 0 and 1, respectively (see Fig. 1**D**). Hence a quasi-equilibrium considered in the following: each of these two birds locates at the best relative longitudinal position (*x*^∗^) to the front neighbor that maximizes the wake benefit from the front neighbor, same as in [16, 24, 28, 34–36, 45]. However, this quasi-equilibrium has a trend to drift downstream due to the wake benefit of bird 1 from bird 2 (*f* (*p*_12_)). Although this benefit *f* (*p*_12_) is much smaller than the wake benefit of bird 1 from the front neighbor (*f* (*p*_10_)), the former increases more than the latter decreases when bird 1 moves backward. This implies that near equilibrium, bird 1 would like to move backward (downstream) as that would increase its wake benefit *f*_1_(*p*) = *f* (*p*_10_) + *f* (*p*_12_). Meanwhile since bird 2 has no back neighbor, it would like to maintain the relative longitudinal position to bird 1 that maximizes its wake benefit *f*_2_(*p*). In cycle, bird 1’s backward motion would “push” bird 2 back, which in return attracts bird 1 to move backward again. Hence the almost equilibrium cannot be maintained. This explanation can be extended to the case with more than 3 birds, in particular to the last two followers, since from Fig. 1**D** if the distance of each bird to its front neighbor is around |*y*^∗^|, it gets the wake benefit mostly from neighbors.

**Fig 4.**
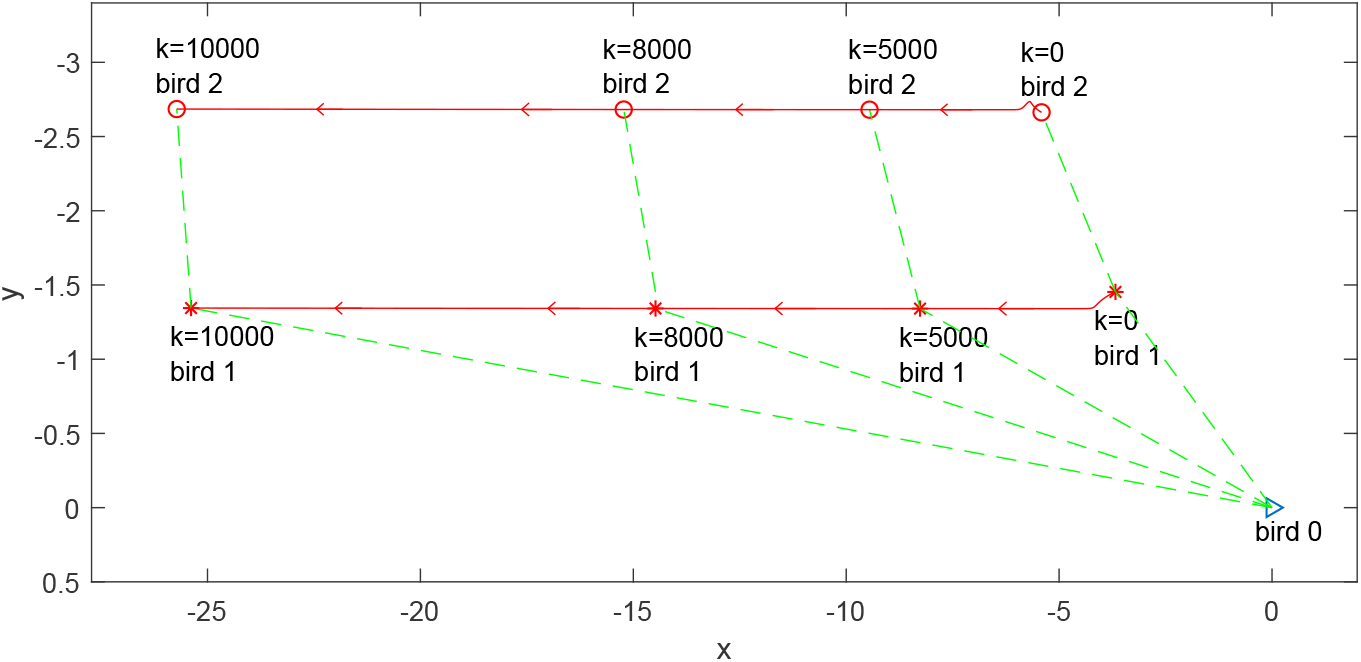
The position history of two birds in the egoistic equilibrium search based on the gradient algorithm. The parameters for the bird size and weight, and the wake model are the same as those in Fig 1. The red lines represent the position of two birds at the different iteration *k* of the gradient algorithm. The arrows on the two lines denotes the direction of the gradient of each bird’s wake benefit.

As for the cooperative case, there does exist a cooperative equilibrium, where the *y* coordinate of bird 1 and 2 is almost *y*^∗^ and 2*y*^∗^, respectively, and the *x* coordinate of both birds equal to zero. The corresponding configuration are birds equally distributed on a laterally extended line, where birds longitudinal positions are the same and the lateral distance of each two neighboring birds equals |*y*^∗^|. This can be regarded as a large wingspan bird. This confirms the observations that: 1) When *x* = 0, the wake benefit *f* (*x, y*) attains the maximum at *y*^∗^ on the negative half *y* axis, as shown in Fig 1**C**; 2) When the lateral positions of birds 1 and 2 are fixed to *y*_1_ = *y*^∗^ and *y*_2_ = 2*y*^∗^, the total wake benefit *J* reaches the maximum at *x*_1_ = *x*_2_ = 0, as shown in Fig. 5. The second observation can be explained if we consider the Munk’s stagger theorem [32], which states that “a collection of lifting surfaces (birds) may be translated in the streamwise direction (*x* direction) without affecting the total induced drag of the system (flock) as long as the circulation Γ of every wing (or lift) is unchanged.” The induced drag of each bird can be decomposed into the self-induced drag and the mutually induced drag [31, 32], where only the latter relates to the net vertical air velocities induced by other birds wakes. The mutually induced drag of each bird is proportional to its wake benefit [31]. Hence one can also conclude that as long as the vortex circulation Γ of every wing is unchanged, the total wake benefit of the flock does not change if birds slide along the streamwise direction. This conclusion holds for constant circulation. However, to count the slow dissipation of wake [44, 45], our wake model assumes that the radius of the trailing vortex increases slowly along the streamwise direction, see Equation (1) in Methods. Although, the real circulation Γ is still constant, this is equivalent to introduce a virtual vortex circulation modulated by a space dependent term that decreases slowly as the longitudinal distance between two birds (namely |*x*|) increases. Due to this decay, the total wake benefit reduces as birds longitudinal distances increases. This shows that *J* achieves the maximum if all the birds longitudinal distances are zero, namely, the second observation. To this end, we point out that the cooperative equilibrium found is not realistic since the distance of every neighboring birds is |*y*^∗^| ≈ 0.893 wingspan, implying the collision of birds. Hence, this shows that if birds collaborate toward the maximization of the total wake benefit of the group, no typically observed formation shape can emerge.

**Fig 5.**
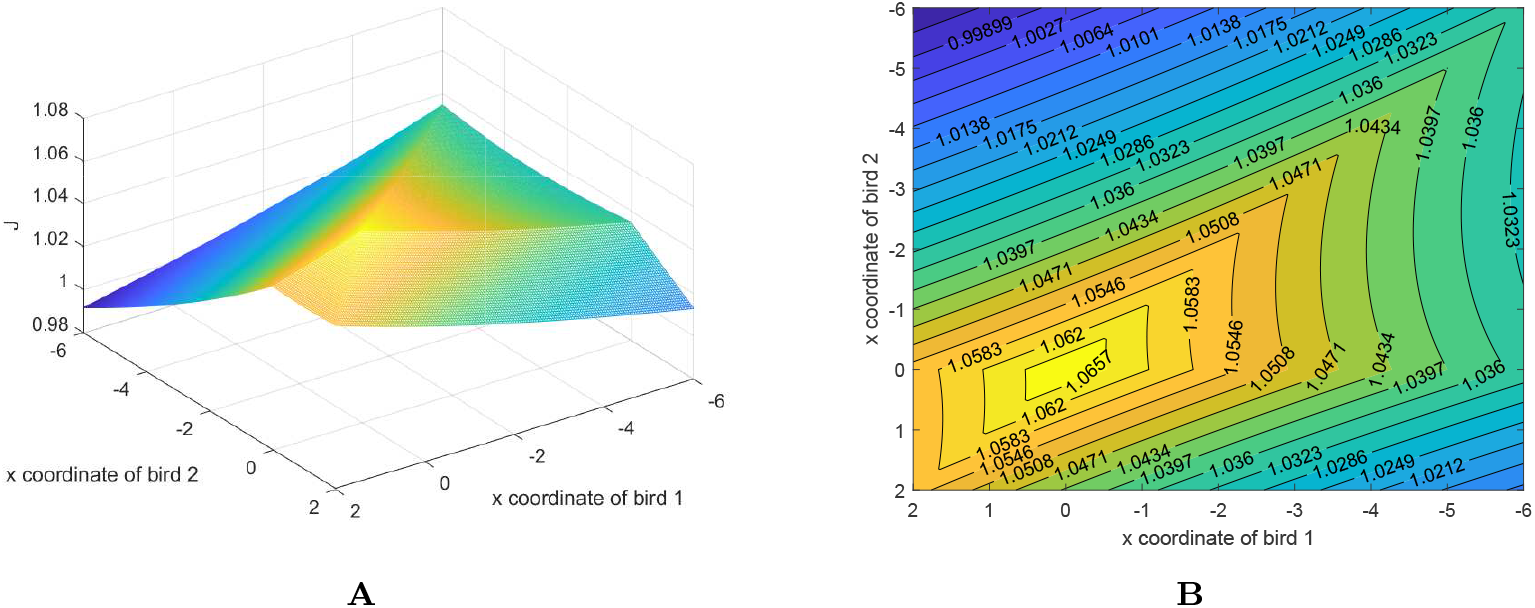
The total wake benefit *J* when *y*_1_ = *y*^∗^ and *y*_2_ = 2*y*^∗^. **A**: The 3D view of *J*. **B**: The contour of *J*.

### Collision avoidance and vision comfort

Let us now waive the hypothesis that birds only optimize wake benefit. Our computations show that echelon formations can emerge if collision avoidance or/and vision comfort keeping are also considered. Hence, these two factors may play important roles in the emergence of the echelon formation of birds in reality. However, their roles are dissimilar for selfish and cooperative birds.

If birds cooperate to optimize the total wake benefit and try to avoid colliding with each other, then there exists one cooperative equilibrium, whose corresponding formation configuration for a flock with 1 leader and 10 followers is demonstrated in Fig 6**A**. Hence being aware of collision risk contributes to the echelon formation emergence. This conclusion can be understood for general collision avoidance models. As shown in the previous subsection where collision avoidance is excluded, to maximize the group wake benefit, the lateral distance of neighboring birds should be around |*y*^∗^| and birds should be cohesive longitudinally. The forbidden region around each bird induced by collision avoidance enforces a non-zero longitudinal separation between neighboring birds when their lateral distance is around |*y*^∗^|. Hence each follower cannot be too close to its front neighbor longitudinally, although that would increase the group’s wake benefit. Indeed, the computation shows that the conclusion is robust to the choice of artificial barrier functions, since the cooperative equilibria for two different artificial barrier models (See Method) exist and the positions of birds corresponding to both equilibria are almost the same. As for selfish birds, avoiding collision with others does not promote the forming of the echelon formation, as no egoistic equilibrium appears in the simulation. It is found that the followers continuously drift back along the steady streamwise direction, same as in the previous subsection.

**Fig 6.**
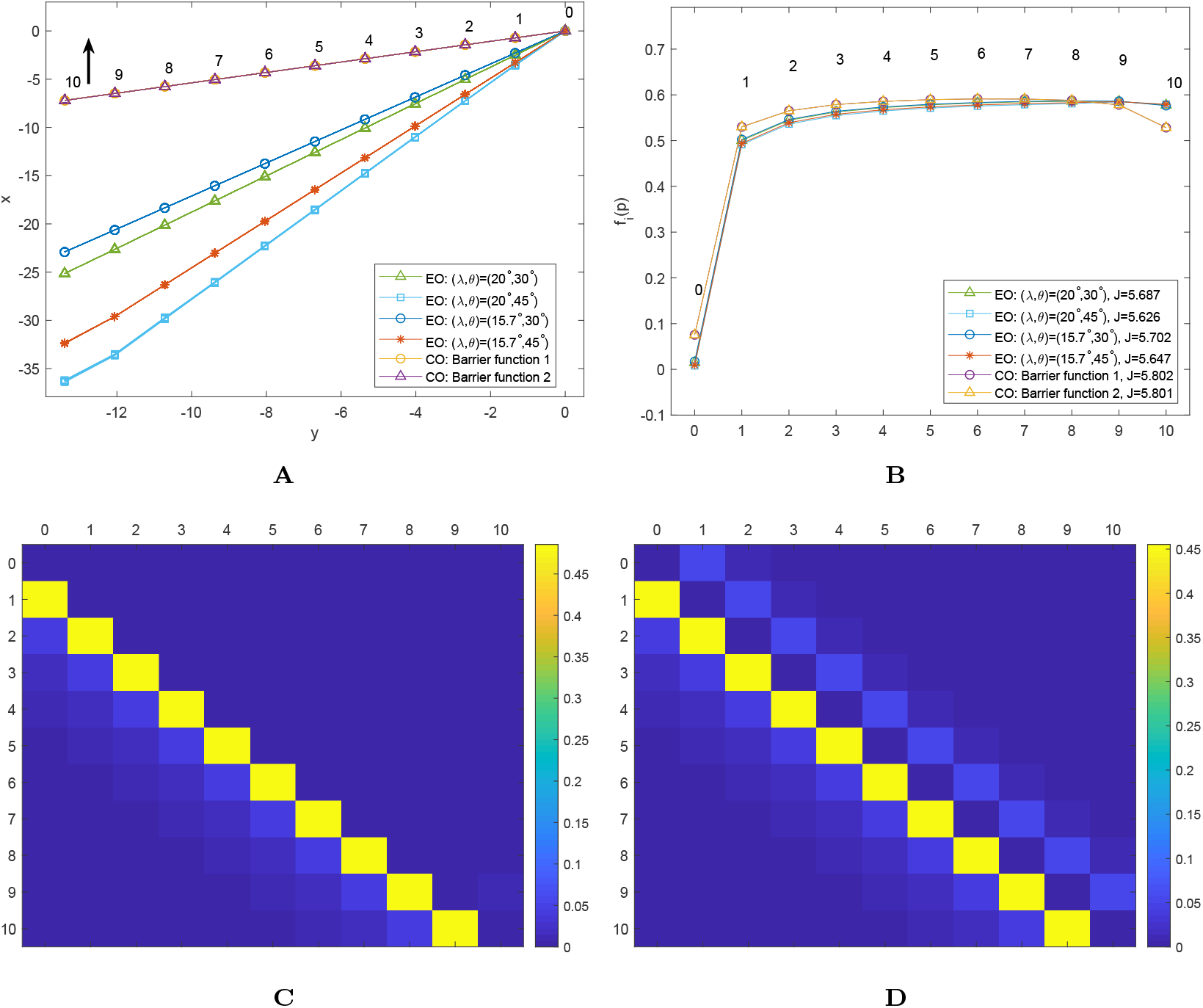
10 + 1 birds formation corresponding to the egoistic equilibrium and cooperative equilibrium. The egoist equilibrium is obtained by the gradient method, including also the gradient of the vision comfort. While seeking the cooperative equilibrium, the gradient method includes the gradient of the wake benefit and the barrier function. We try two different barrier models and four cases of the vision angle parameters ((*λ, θ*) = (20°, 30°), (20°, 45°), (15.7°, 30°), (17.5°, 45°)). Here, the vision comfort angle *θ* is selected arbitrarily, and the front binocular angle *λ* = 15.7° for the last two pairs of vision angles is based on the measurement from Canada geese [37]. For each vision angle pair and barrier model, the equilibrium is searched for 50 different random initial birds positions. See Method for details. **A**: Formation configuration for the two equilibria. EO and CO represent the egoistic equilibrium and cooperative equilibrium, respectively. **B**: The wake benefit value for each bird. **C**: The benefit of each bird *i* in the row induced by wake vortices of bird *j* in the column at the egoistic equilibrium with (*λ, θ*) = (15.7°, 45°). **D**: The benefit of each bird *i* in the row induced by wake vortices of bird *j* in the column at the cooperative equilibrium with barrier model 1.

On the other hand, keeping vision comfort plays a part in forming the echelon formation for selfish birds, since there exists one egoistic equilibrium if birds simultaneously optimize the wake benefit and the vision comfort. Examples are illustrated for a flock with 1 leader and 10 followers and four pairs of vision angles. Computation shows that the egoist equilibrium exists for all the four pairs, with the resulting bird formation given in Fig. 6**A**. From this figure, at the egoistic equilibrium, the lateral distance of the follower to the front neighbor within and across all the four cases of (*λ, θ*) pair are almost the same. However, this is not the case for the longitudinal distance of neighboring birds. A wide vision comfort zone (large *θ*) and a wide front binocular vision (large *λ*) can both lead to the formation, where birds are much separated in the streamwise direction. The pair (*λ, θ*) = (20°, 30°) and (*λ, θ*) = (15.7°, 30°) lead to a shallow formation angle that is spanned by the line of birds positions and the horizontal line, while the pair (*λ, θ*) = (20°, 45°) generates a slightly distorted flocking configuration, where the longitudinal distance of bird 10 to 9 is less than that of other followers to their front neighbors. This is still reasonable since in the real birds formation, the longitudinal distance of neighboring birds has a large variation [35]. From this, we conclude that the echelon formation can emerge if each bird optimizes its own wake benefit and maintains the vision comfort together. Moreover, a closer look at S1 Table shows that the gradient of the individual wake benefit and the vision comfort of each bird at the egoistic equilibrium are small. This indicates that little effort of birds in reducing vision discomfort can help the formation emerge. Conversely, minimizing vision discomfort does not assist cooperative birds in achieving the formation, as no cooperative equilibrium related to the line formation exists. We only find the same equilibrium as in the previous Subsection, where all followers are in alignment with the leader at the same longitudinal position and the distance of neighboring birds is less than the wingspan. This follows from the fact that each front bird already positions itself within the vision comfort zone of its back neighbor at this equilibrium. However, as mentioned before, this equilibrium is not physically feasible as birds would collide.

When collision avoidance and vision comfort are both combined with energy benefit optimization, the echelon formation emerges for both cooperative and selfish birds: both egoistic and cooperative equilibria exist. The egoistic equilibria for the four cases of (*λ, θ*) are identical to those obtained if collision avoidance is excluded. Similarly, the cooperative equilibria for barrier functions 1 and 2 are the same as those obtained if vision comfort is not included. Table 1 summarizes the factors that affect the appearance of equilibria. At all the equilibria, the lateral distance of neighboring birds is almost the same (closely around 0.893 wingspan, same as in [22, 28, 31]), but the longitudinal distance of neighboring birds varies much. Even so, the total wake benefit *J* (excluding the value of the barrier potential and the vision comfort) of the egoistic equilibrium for all four cases of (*λ, θ*) does not differ too much from that of the cooperative equilibria, see Fig. 6**B**). A wider vision comfort zone and a more front-biased binocular vision cause the total wake benefit of the flock to decrease more. The small variation of the total wake benefit *J* even when the longitudinal position of birds varies in a large range, is due to the slow decay of the wake benefit function *f* (*p*) on the line *y* = *y*^∗^ along the streamwise direction.

**Table 1.** Collision avoidance and vision comfort vs equilibria existence.

|  | CA | VC | Both | Neither |
| --- | --- | --- | --- | --- |
| Egoistic equilibrium | × | ✓ | ✓ | × |
| Cooperative equilibrium | ✓ | × | ✓ | × |

The table shows whether the equilibria exist under the factor of collision avoidance or/and vision comfort. The mark ✓ (×) represents that the equilibrium exists (not exists). "CA" and "VC" stand for collision avoidance and vision comfort, respectively.

As mentioned in Model and also well recognized in [30, 31]. Fig 6**B** shows that the leader bird gains wake benefit from other birds, even though it is much smaller than the wake benefit of followers. The wake benefit of every bird, except the last one, at the egoistic equilibrium is no greater than that at the cooperative equilibrium. This is because, in the case of selfish birds, no follower cares about others birds. The decrease of the wake benefit of the last bird for the cooperative equilibrium is more tricky, and we try to explain it hereafter. Note that the last bird can receive the wake benefit only from front birds. Based on Fig. 1**C** and 1**D** and birds relative positions at the equilibria in Fig. 1**A**, the benefit of the last bird induced from the closed front bird (the second to last bird) for the cooperative equilibrium is less than that for the Nash equilibrium, since the last bird at the cooperative equilibrium is located longitudinally before the position where the wake benefit it receives from the second to last bird reaches the maximum. Also, the benefit of the last follower induced from other front birds seem the same for both selfish and cooperative cases. Hence, the last bird at the cooperative equilibrium receives less wake benefit than at the Nash equilibrium. Next, from Fig. 6**C** and 6**D**, each bird seems affected by four upstream birds (when present), but only one (selfish case) or two (cooperative case) downstream birds. For every follower, the largest contributor for both the egoistic equilibrium and the cooperative optimum is the neighbor immediately upstream. The situation differs for the second largest contributor: it is the second nearest upstream bird (when present) and the downstream neighbor for the selfish case and the cooperative case, respectively. This indicates that a bird gets a larger wake benefit from its downstream neighbor when birds are cooperative. As for the leader, the downstream neighbor is the largest contributor of wake benefit in the cooperative birds case.

Finally, other equilibria where the flock is split into sub-flocks could exist. In fact, when birds are selfish, if we artificially select birds’ positions such that there exists a bird *i* that is far away from its front neighbor in the lateral direction, e.g., 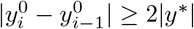, then the leader and the first *i* − 1 followers would form a sub-group, while the last *n − i* + 1 birds would form another subgroup. Within each sub-group, the relative lateral position of each bird to its front neighbor is *y*^∗^. The flock split is caused by the vision discomfort mitigation and the slowing change and small value of the wake benefit *f* (*p*) when |*y*| ≥ 2 |*y*^∗^| : due to the latter reason, birds *i, i* + 1, …, *n* nearly receive no wake benefit from birds 0, 1, …, *i* − 1 and their incentive to get closer to *i* − 1 laterally is very weak. Hence bird *i, i* + 1, …, *n* can be considered as a free moving group, with *i* the leader, and will drift back along the stream direction, since from Fig 1**D** the leader gains more wake benefit if it is closer to the back neighbor. However, to keep the front neighbor *i* − 1 close to bird *i*’s vision comfort zone, bird *i* cannot move back too much, and would stop somewhere, making the back birds also stop. The above analysis shows that line formations can easily break apart if any two neighboring birds are not close laterally, which can be caused by turbulence or some wind shear. Hence it would be hard for large scale flocks to keep the formation intact. This also poses a problem for birds to initiate a line formation and implies that birds have a certain knowledge about the optimal position relative to the front birds, possibly after long time learning and evolution.

### The effect of empathy on energy saving

We now discuss an intermediate case where birds are empathetic and concerned about the energy expenditure of neighbors. Empathy has been observed in some birds, e.g, the social interaction of bystanding greylag geese involved with paired partners within the same family [47]. Furthermore, [48] showed that paired birds tying together and interacting with fewer close birds may save more energy for the pairs. One way to interpret this is that birds show empathy to other close-related birds and care about their performance during flight. Though this is only found in bird clustering, we may expect this also happens for formations of migratory birds of the same family. Note that not all migratory formations are composed of members of the same family, hence, birds empathy is unlikely the key reason behind line formation. This also explains why we consider empathy separately from the selfish and cooperative birds assumptions. We model the empathy by assuming that each follower maximizes its wake benefit plus the weighted wake benefit of the front and back neighbors, with the weight *h* ∈ [0, 1]. Increasing *h* implies that birds care more about their neighbors. By restricting *h* ≤ 1, we assume that each follower cares about neighbors no more than itself. Followers maximizing the mixed benefits form another non-cooperative game, whose Nash equilibrium is named as the empathetic equilibrium (EmE2) in this paper. We also consider the case where each bird only shows empathy to its front neighbor, and thereby maximizes its wake benefit and the weighted wake benefit of its front neighbor. Similarly, this leads to another game whose empathetic equilibrium (EmE1) may exist. We note that [47, 48] point out the possibility of empathy with more birds in the formation, which would lead to more complex models. Nevertheless our model already accounts for empathy towards agents with which the main aerodynamic interactions take place, and should therefore account for the main effects.

We compute the empathetic equilibria also by the gradient method, in which the gradient of the barrier function *B*_*c*_ and the vision comfort function *B*_*v*_ are combined with the gradient of the mixed wake benefit. Note that our model of empathy assumes that all the birds have the same degree of empathy and birds can access their neighbors’ wake benefits and their gradients with sufficient accuracy. One should bear in mind that birds can never “feel” or measure the wake benefit of neighbors. They may get these information only by communication, e. g., honk, through which the exchanged information are always coarse and noisy. However, we stress again that our focus is the equilibria, not the emergence of birds empathy and the way that birds seek the equilibria. The computations of the empathetic equilibria have not led to the observation-matching echelon formation when birds only maximize their mixed wake benefit. In the following we only focus on the situation where vision discomfort mitigation and collision avoidance also influence birds behavior.

We observed that when *h* is close to 1, multiple equilibria appear for both empathy models, see EmE2 with *h* = 1 in Fig 7**A** and EmE1 with *h* = 0.8, 0.9, 1 in Fig 7**C**. Also there is a trend that as *h* increases, EmE2 and EmE1 shift from the egoistic equilibrium towards the cooperative equilibrium obtained in the previous subsection, and the longitudinal distance of some birds to their front neighbors decreases, even though this is not obvious for EmE2, see Fig 8 and S1. In contrast, the lateral distance of neighboring birds is almost the same as that of the egoistic equilibrium or the cooperative equilibrium. For the situation where birds are empathetic to both front and back neighbors, the further down the bird is in the formation (except the last bird), the higher its wake benefit will be, see Fig. 7**B**. This also holds for the situation where birds only care about the front neighbors with little empathy (e.g. *f*_*i*_(*p*) for EmE1 with *h* = 0.1, 0.2…, 0.9), but is not true when *h* = 1, where the bird could get more wake benefit if it locates in the middle of the formation. In addition, for both empathy models, if birds care more about their neighbors, the whole group is more energy efficient, since the difference between the total benefit at the empathetic equilibria and that at the cooperative equilibrium is smaller, see Fig. 8. Interestingly, this figure also shows that the energy efficiency losses for all EmE1s are smaller than that for all EmE2s when *h* ≥ 0.6. This means that the flock with followers being only empathetic to their front neighbors get more wake benefit or saves more energy than the flock with followers being empathetic to both the front and back neighbors. All in all, Fig. 8 shows that if empathy does exist in close-related birds, then these birds flying together could save more total energy than if they were purely selfish.

**Fig 7.**
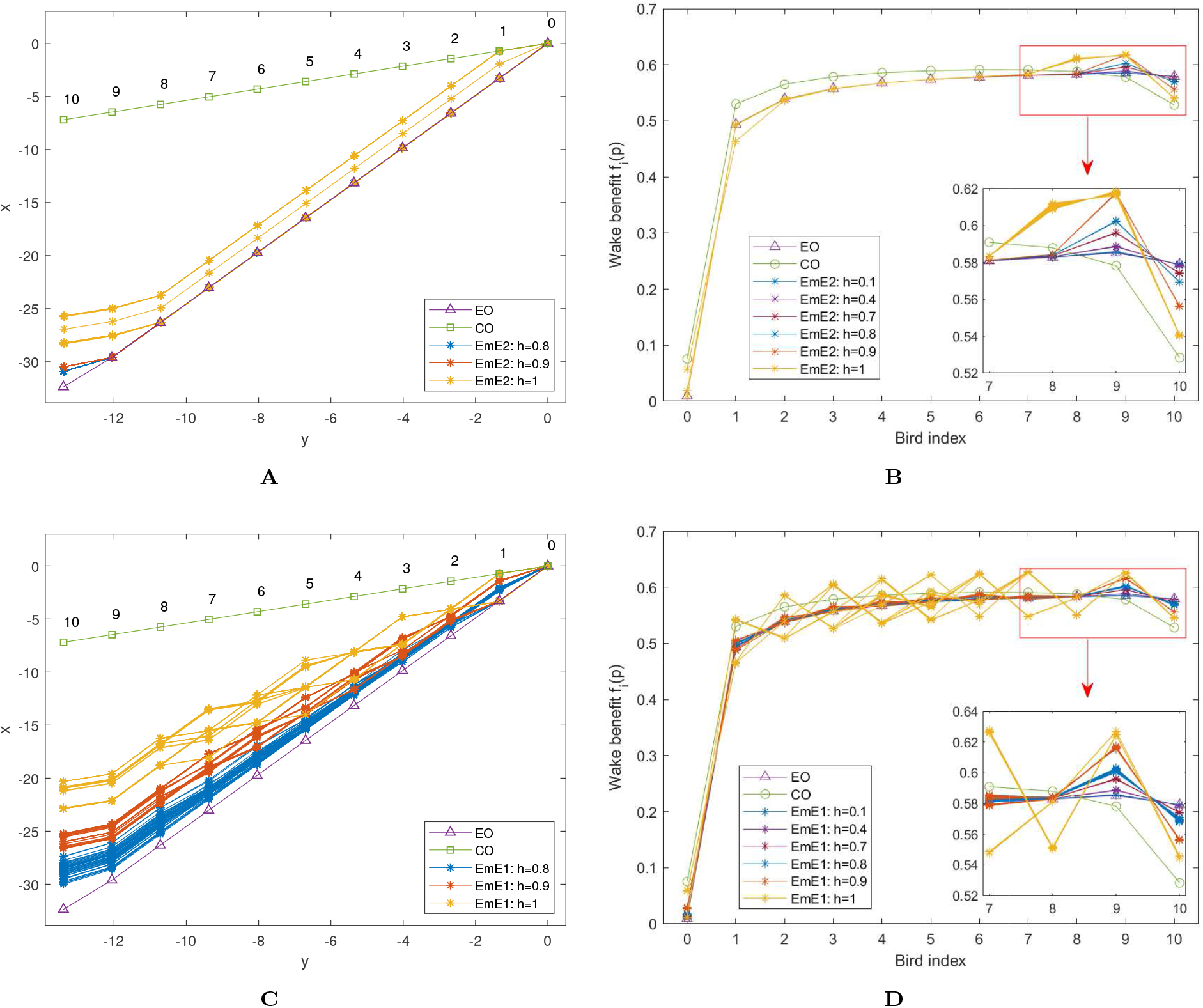
Formation for empathy flocks with 1 leader and 10 followers. In the empathetic equilibria search, we select the barrier function 1 and the vision angles (*λ, θ*) = (15.7°, 45°). For each empathy degree, the equilibrium is searched for 50 different random initial birds positions and multiple equilibria could be obtained for each empathy degree. The results are compared with those of the cooperative equilibrium (CO) with the same barrier function and the egoistic equilibrium (EO) with the same vision angles, obtained in the previous section. See S1. Fig. for the equilibria when birds show empathy to both front and back neighbors (EmE2) and only to the front neighbors (EmE1), respectively, with *h* = 0.1, …, 0.7. **A**: Formation at the EmE2 with *h* = 0.8, 0.9, 1. **B**: Wake benefit of each bird at the EmE2. **C**: Formation at the EmE2 for *h* = 0.8, 0.9, 1. **D**: Wake benefit of each bird at the EmE1.

**Fig 8.**
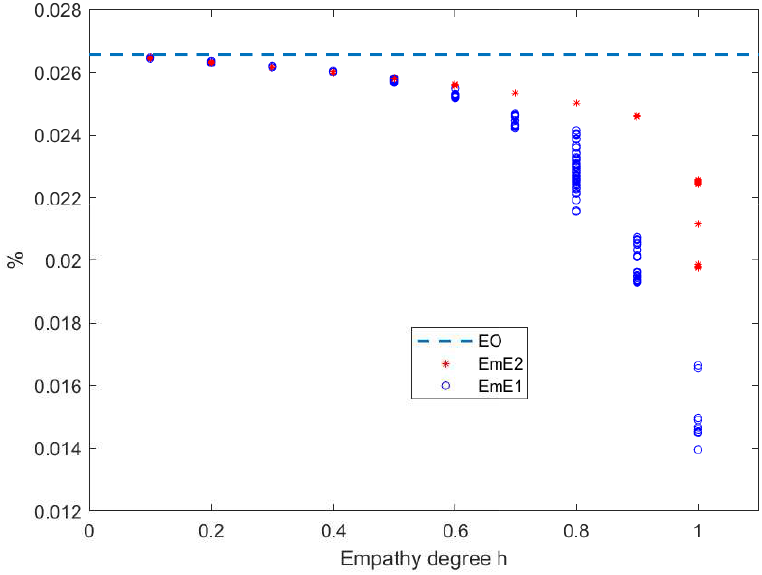
The energy effiency loss of the flock at the empathetic equilibria in Fig. 7. The energy effiency loss is defined as 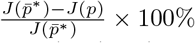, where 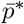 is the cooperative equilibrium obtained for the case with (*λ, θ*) = (15.7°, 45°) in the previous subsection and *p* is the birds positions corresponding to a formation shape. The dashed line represents the energy loss of the group at the egoistic equilibrium.

## Discussion

Even though energy saving has been known to play an important role in the formation flight of large migration birds, the question of how birds exploit this benefit is still open. Using a modified fixed-wing model for bird flying, we revisit the the emergence of the common migration formation, by testing whether any stationary echelon formation shape can be reconstructed numerically when birds are selfish or cooperative somehow in maximizing the wake benefits. We demonstrated that the hypothesis of birds purely optimizing their aerodynamic wake benefits is not sufficient to produce a formation that is similar to the daily-observed ones, for both cooperative and selfish birds. On the other hand, collision avoidance and vision discomfort mitigation can assist the flock in creating the line formation. Collision avoidance does help cooperative birds, which optimize the total wake benefit of the entire flock, but not selfish ones, which maximize their own wake benefits, toward forming an echelon formation. As a contrast, mitigating vision discomfort or enhancing vision could play a part in creating echelon formations for selfish birds, but not cooperative birds. Moreover, if reducing collision risk and maintaining vision comfort are assumed to simultaneously play roles, then echelon formations appear for both cooperative and selfish birds, as well as for birds that are empathetic and maximize their own and the neighbors’ wake benefits. Hence, we conclude that the motivation of birds during long-term flights is to optimize the wake benefit (or reversely reduce energy consumption). However, the well-known formation shape depends on non-aerodynamic factors: the collision avoidance and/or the vision constraint. Across and between the constructed formations for all the cases of self, cooperative and empathetic birds, the lateral distance between neighboring birds is almost the same, but the longitudinal distance between neighboring birds differs much.

In addition, we found that the total wake benefit of the flock for the formation obtained by considering non-aerodynamic factors is most maximized for cooperative birds, followed by empathetic birds, and then selfish birds. However, the differences among the three cases are quite small, even though the longitudinal position of birds across the three situations differ much. Hence, compared to the cooperative situation, the energy efficiency for migration flocks following partially cooperative, or even purely selfish strategies is very high. This is due to the fact that in the three situations, each follower gets the most wake benefit from the front neighbor, which does not change much when the follower moves along the streamwise direction. It happens because the wake benefit generated by the upwash behind a flying bird peaks along two lines that extend in the streamwise direction, and varies vastly around these two lines in the lateral direction but decays very slowly along the streamwise direction. This also causes the absence of line formation emergence for selfish birds that purely optimize wake benefits. Due to the feature of the wake benefit, if the bird moves backward along the streamwise direction, the wake benefit increment due to the back neighbor is larger than the loss of wake benefit due to the front neighbor. This, along with the fact that the last bird in the flock achieves the maximal wake benefit when it keeps an almost constant longitudinal distance to the front neighbor, leads to a cyclic dilemma for the last two followers: the second to last follower’s backward motion increases it own benefit but reduces that of the last follower, while the backward motion of the last follower increases its own benefit but reduces the second last follower’s.

Although energy-guided motions with either collision risk reduction or vision discomfort mitigation can lead to the (approximate) line formation for the three situations of birds interests, inferring which behavior is really followed by birds and which non-aerodynamic factors matter to the line formation emergence in reality is difficult. It is easy to admit that collision avoidance is critical in formation flight, while the vision comfort may only be an artificial factor and how birds perceive it and take action to maintain it is unclear. Considering these, one may conjecture that birds are indeed cooperative since the echelon formation can be reconstructed for cooperative birds excluding the vision comfort factor. However, the cooperative birds assumption requires all the birds to optimize the total wake benefit of the entire flock. Obviously, the complexity of the wake prevents birds to know the mathematical form of the total wake benefit. Hence, to optimize the total benefit, each bird would need to have the ability to access other birds’ positions information and their perceived wake benefits. Whether this is possible and the precise way in which it would be achieved remain open questions. Birds might also need to communicate with all other birds to get the information for optimization. However, as mentioned before, the communication among birds is probably very coarse due to wind noise and birds may not have sufficient ability to process information. Considering these, it is undoubtedly a difficult task for birds to cooperatively find the positions that most reduce the energy cost of the group the most. For empathetic birds, each follower also needs to obtain the wake benefit of its neighbors’, hence the problem of coarse communication again, which could prevent the current simulation result to hold. In contrast, selfish birds only need to perceive the gradient of their own wake benefits, hence we conjecture that the assumption of selfish birds is closer to the reality when the flock size is large. Another argument for this conclusion is that although birds behave selfishly, the energy efficiency of the flock does not deteriorate much. Hence, birds may lack the motivation to switch or learn to behave collaboratively. The reconstruction of the echelon formation for selfish birds relies on the assumptions that vision comfort zones exist for migrating birds and birds eyes are relative immobilized during the long-time migration flight. We have not made experiments to justify these assumptions and quantify the vision comfort model. However, we anticipate they are qualitatively correct based on the experimental research in [37, 39, 40].

The failure in reproducing the echelon formation without considering the non-aerodynamic factors may also be caused by the simplicity of the current models. We have indeed regarded each bird as a fixed-wing, and only take the averaged vertical aerial velocity generated by other birds as the wake benefit. A faithful model would account for the bird’s kinematics, flapping gait and the resulting aerodynamics [51]. Additionally, the wake of a flapping bird is unsteady by nature and it will more likely lead to a varying benefit or other stabilizing or destabilizing effects on a follower’s position.

## Methods

### Wake model and aerodynamic benefit function

Consider a leading bird with weight *W* and wingspan *b* flying with speed *U* in the *xy* plane along the *x* direction and assume that the air density is *ρ*. If approximating birds by fixed-wings, the most widely used model to represent the wake of the leading bird is the horseshoe model, in which the wake consists of a finite bound vortex and two trailing vortices, The forming of these vortices is briefly explained as follows and can be found in [32, 50].

To support the bird weight, a difference of the air pressure between the lower and upper surfaces of the wing should be sustained. Since this difference cannot be maintained at the wingtip, where no surface isolates the top and bottom air, the air with higher pressure from the inboard and bottom of the wing moves to the low-pressure wing outboard and wing top. This causes the air at the wingtips to roll up into two semi-infinite trailing vortices with the the same circulation 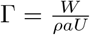 and opposite directions. The two trailing vortices start just inboard of the wingtips and extend downstream. Moreover, the air pressure difference is usually explained by the acceleration and deceleration of the airflow above and below the wing, respectively, relative to the free stream. The difference in the airflow speed above and below the wing can be regarded as a circulatory flow around the wingspan, attached to the free stream. This circulatory flow is the finite bounded vortex, with length 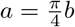 and circulation Γ.

However, as mentioned in Introduction, the trailing vortex strength decays slowly as it extends downstream [44, 45]. To account this decay, we used a modified horseshoe model, given hereafter. Suppose the leading bird is at the origin [0 0]^⊤^, the vertical airflow velocity *v*(*p*) generated by the vortices at *p* = [*x y*]^⊤^ can be given as

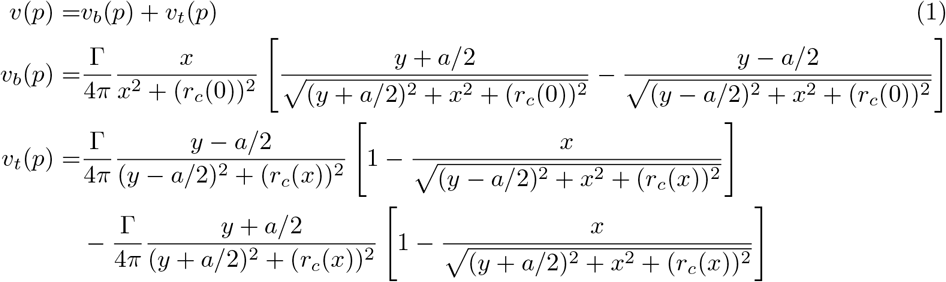

where *v*_*b*_ and *v*_*t*_ are the vertical velocities induced by the bound vortex and the trailing vortices, respectively, 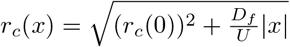, *r*_*c*_(0) is the vortex core radius at *x* = 0 and taken as 0.02*b* in this paper, and *D*_*f*_ is the diffusion term. The introducing of *D*_*f*_ allows the vortex core to expand as it gets away from the wing along the streamwise direction. This modification enable us to incorporate the decay phenomenon of the trailing vortex circulation. We select *D*_*f*_ = 5.25 × ^−5^*Ub* such that *r*_*c*_(*x*) increases from 0.02*b* to approximate 0.05*b* when |*x*| varies from 0 to 40*b*, which is fairly realistic according to the empirical data [49].

Then consider a following bird with the same size and weight as the leader. Let the center of the follower be *p* = [*x y*]^⊤^. We neglect the momentum induced by the vertical airspeed, and characterize the wake benefit of the following bird generated by the leading bird as follows,

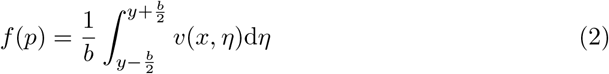

In the simulation of this paper, all the birds are assumed to have the same size and weight. The parameters in the wake model are set as *b* = 1.5m, *W* = 36.75N, *U* = 18m/s^2^, *r*_*c*_(0) = 0.02*b* and *ρ* = 1.124kg/m^3^,

### Egoistic equilibrium and cooperative equilibrium

For bird *i* = 0, 1, …, *n* with position *p*_*i*_ ∈ ℝ^2^, its net wake benefit is assumed to be

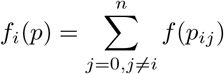

where *p*_*ij*_ = *p*_*i*_ − *p*_*j*_ is the relative position of bird *i* to bird *j*, and *f* (*p*_*ij*_) is the wake benefit of bird *i* induced by the wake generated by bird *j*.

When birds are selfish and only maximize their own wake benefits, we denote the egoistic (Nash) equilibrium by 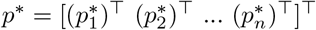, where 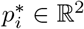 is the position of bird *i* at the equilibrium. According to the definition of Nash equilibrium, if *p*^∗^ exists, it should satisfy

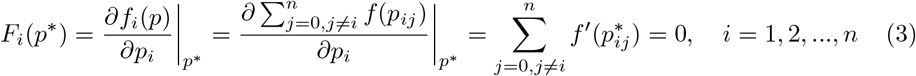

where 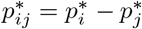 and *f* ^′^(·) denotes the first derivative of *f* (·).

The cooperative equilibrium is the point at which the following total wake benefit function is maximized,

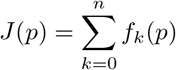

Denote by 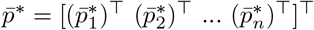, with 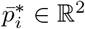 for *i* = 1, 2, …, *n*, the cooperative equilibrium. Since *J* is differentiable, 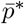 should satisfy the following condition

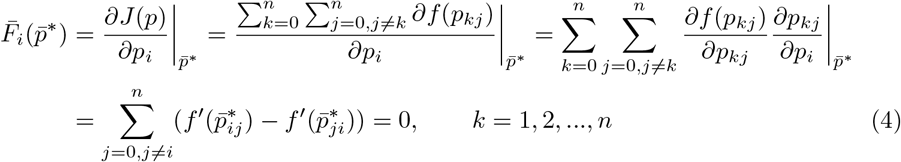

where 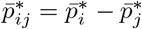.

### Invariance of scaled equilibria under parameters variation

One can show that the equilibria defined in the previous subsection is scale-invariant, in the sense that the scaled versions of the equilibrium remain unchanged when one modifies parameters such as the air density *ρ*, bird weight *W*, wingspan *b* and velocity *U*. Such modifications have thus no effect on the general shape of the formation.

To see this, we first note that multiplying the wake benefit function *f* (*p*) in (2) by the non-argument constant 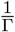 does not change the equilibria. Now let 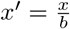 and 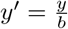 and 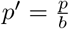 be the scaled position variables, then

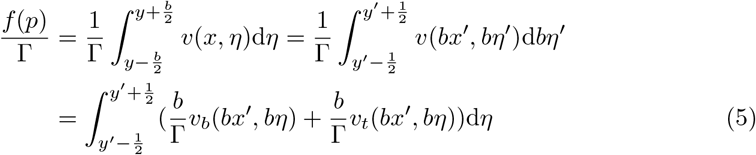

Recalling 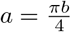, *r*_*c*_(0) = *c*_1_*b*, with *c*_1_ = 0.02, *D*_*f*_ = *c*_2_*Ub*, with *c*_2_ = 5.025 × 10^−5^, and noticing 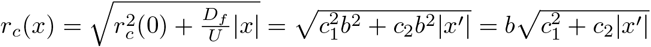, the two terms in the last integration of the above equation can be written as

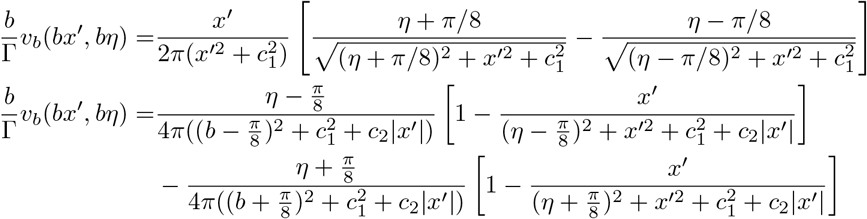

From these, we know that 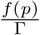 does not contain the parameters mentioned at the beginning of the subsection. This shows that if birds purely maximize wake benefits, the existence of equilibria for the benefit maximization game and total benefit optimization does not depend on these parameters. Moreover, the equilibria (if they exist) normalized by the wingspan do not change as these parameters vary. Finally, we note that if collision avoidance or vision comfort are also considered, the equilibria would depend on the wingspan and bill-to-tail distance of birds, which may vary for different bird species.

### Barrier function

As mentioned in Model, we model each bird as an ellipse, with the semi-major axis *b*_*l*_ and the semi-minor axis *b*_*s*_, where *b*_*l*_ and *b*_*s*_ depend on the size of birds. It is easy to know that the space occupied by any bird with center *p*_*c*_ = [*x*_*c*_ *y*_*c*_]^⊤^ can be given by the following ellipse,

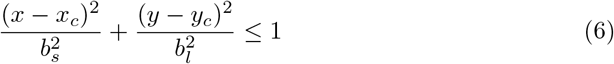

Now consider a bird *j* and any other bird *i*, with the relative position to bird *j* being *p*_*ij*_ = [*x*_*ij*_ *y*_*ij*_]^⊤^. To avoid the collision with bird *j*, as shown in Figure 2**B**, the center of bird *i* should satisfy the condition below,

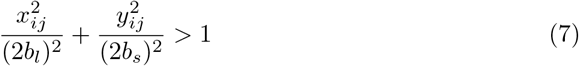

The yellow ellipse in Fig 2**B** that defines the collision alert region of bird *i* with respect to bird *j*.

We use two potential functions modeling the barrier. Let *p*_*ij*_ = [*x*_*ij*_ *y*_*ij*_]^⊤^. Barrier function 1 is given as

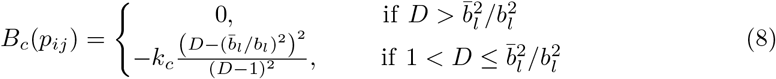

where *k*_*c*_ ≥ 0, 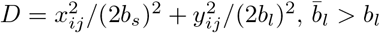 determines the collision alert region.

Barrier function 2 is given as

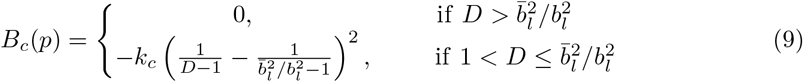

Apparently, for both barrier functions, if bird *i*’s center is outside the collision alert region of bird *j*, namely 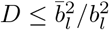, then it considers that it is sufficient far from bird *j*. If *p*_*ij*_ is within the ellipse 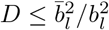 but outside the smaller ellipse *D* ≤ 1, then bird *i* gets a negative value for the barrier function, since its metabolic level may rise due to the pressure of the possible collision. In the simulation, we select *k*_*c*_ = 1, 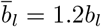 and set parameters as 2*b*_*l*_ = *b* = 1.5m and 2*b*_*s*_ = 0.9m (Canada goose).

### Vision comfort function

We give the vision comfort function *B*_*v*_(*ϕ*) here. Since the left-V formation is considered in this paper, we only focus on birds right eyes, but emphasize that the case for left eyes can be derived in an analogous way. Note that the precise shape for the vision comfort zone and the explicit expression of the function *B*_*v*_(*ϕ*) are open to discuss, but the main effect of maintaining eye comfort remains. Consider any bird *i* and its front neighbor *j* and define the vision angle *ϕ* of bird *j* in bird *i*’s eyes be the angle from the right vision axis of bird *i* to the ray connecting the right eye of bird *i* (the front end of the semi-minor axis of the ellipse) to the center of bird *j*, counterclockwise, see Fig 3**B**. Note that *ϕ* can be negative. Let *s* = tan *ϕ*, then |*ϕ*| ≤ (*>*)*θ* is equivalent to |*s*| ≤ (*>*) |tan *θ*|. Compared to *ϕ, s* is can more easily be obtained from the relative position of bird *i* to bird *j, p*_*ij*_ = [*x*_*ij*_ *y*_*ij*_]^⊤^. As shown in Fig 3**B**, the counterclockwise angle from the *y* axis to the right vision axis of bird *i* equals the front binocular vision angle *λ*. Let *ψ* be the counterclockwise angle from the *y* axis to the ray connecting bird *i*’s right eye to the center of bird *j*. Then there holds *ϕ* = *ψ* − *λ*. Since 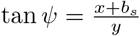 from Fig 3**B**, *s* can be computed by the sum formulas for tangent as

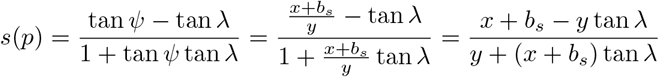

The exact value of *λ* and *θ* depend on the bird. We define the vision comfort function as

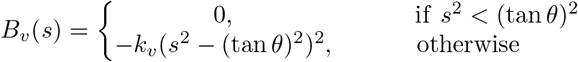

where *k*_*v*_ *>* 0 can be arbitrarily selected and we select *k*_*v*_ = 1 in the simulation. It is easy to see that if |*ϕ*| ∈ *θ, B*_*v*_(*s*) = 0, representing no vision discomfort. While if |*ϕ*| *> θ, B*_*v*_(*s*) *<* 0, meaning that the vision discomfort appears. As 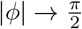 approaches +∞ and the value of function *B*_*v*_(*s*) approaches −∞. However, from the geometry in Fig 3**B**, this cannot happen in the scenario of this paper, since it would imply that either bird *j* locates at the left side of bird *i*, or bird *i* locates at the front of bird *j*. This implies that the possible emergence of the line formation is not enforced by the vision factor as the absolutely dominant. We note that the defined *B*_*v*_(*s*) is indeed a function of the relative position of bird *i* to *j*. Bird *i* adjusts its position to maximize this vision comfort function.

### Gradient based method for testing the equilibria’s existence

To test the existence of the egoistic equilibrium and cooperative equilibrium with or without including the factor of collision avoidance or/and vision comfort, we modify gradient method [41] by incorporating the gradient of the barrier function *B*_*c*_ or/and the vision comfort function *B*_*v*_. In detail, for a flock with one fixed leader and *n* followers, let *p*^0^ ∈ ℝ^2*n*^ be the initial vector of followers’ positions. The iteration for searching the egoistic equilibrium can be given as

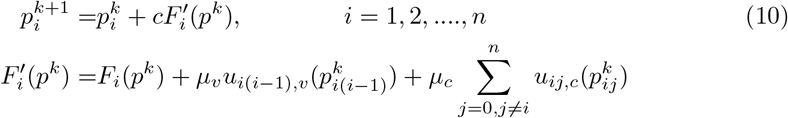

where *F*_*i*_(*p*^*k*^) is given as in (3), 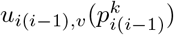 and 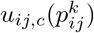 are given as follows,

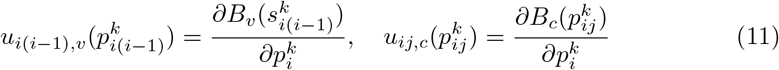

where *s*_*i*(*i*−1)_ = tan *ϕ*_*i*(*i*−1)_, with *ϕ*_*i*(*i*−1)_ denoting the vision angle of bird *i* − 1 in the eye of bird *i*. The variable *μ*_*v*_, *μ*_*c*_ ∈ {0, 1} indicate that whether the collision avoidance or/and maintaining vision comfort are taken into account in the egoistic equilibrium search.

Similarly, the algorithm for search the cooperative equilibrium with or without including the factor of collision avoidance or/and vision comfort can be given as

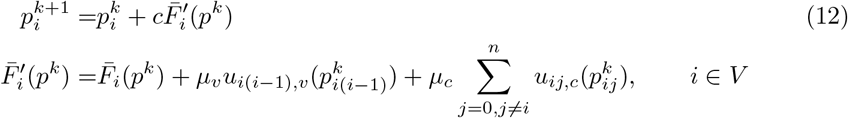

where 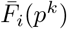 is given as in (4).

If max_*i*=1,2,…*n*_ |*F* ^′^(*p*^*k*^) | ≤ 0.0001 (max_*i*=1,2,…*n*_ |*F* ^′^(*p*^*k*^) | ≤ 0.0001) are satisfied within sufficiently number of iterations, or *p*^*k*^ stays around a point for a large number of iterations, the algorithm (10) ((12)) is considered as converging, otherwise not. We change *μ*_*v*_ and *μ*_*c*_ to see how collision avoidance or/and vision comfort affect the existence of the egoistic equilibrium and cooperative equilibrium, by checking if the search algorithm (10) or (12) for the corresponding cases converges or not. For each different parameters of vision angles and barrier models, we search the egoistic equilibrium and/or cooperative equilibrium for 50 different initial birds positions. The initial birds positions in searching the egoistic equilibrium are chosen such that the longitudinal distance and lateral distance of each follower to its front neighbor are randoms within the interval [0.75, 5.15] and [1.06, 1.66], respectively. For the search of cooperative equilibrium, these two initial distances of each neighboring birds are selected randomly within [0.75, 1.75] and [1.06, 1.66], respectively.

## Supporting information

**S1 Fig.**
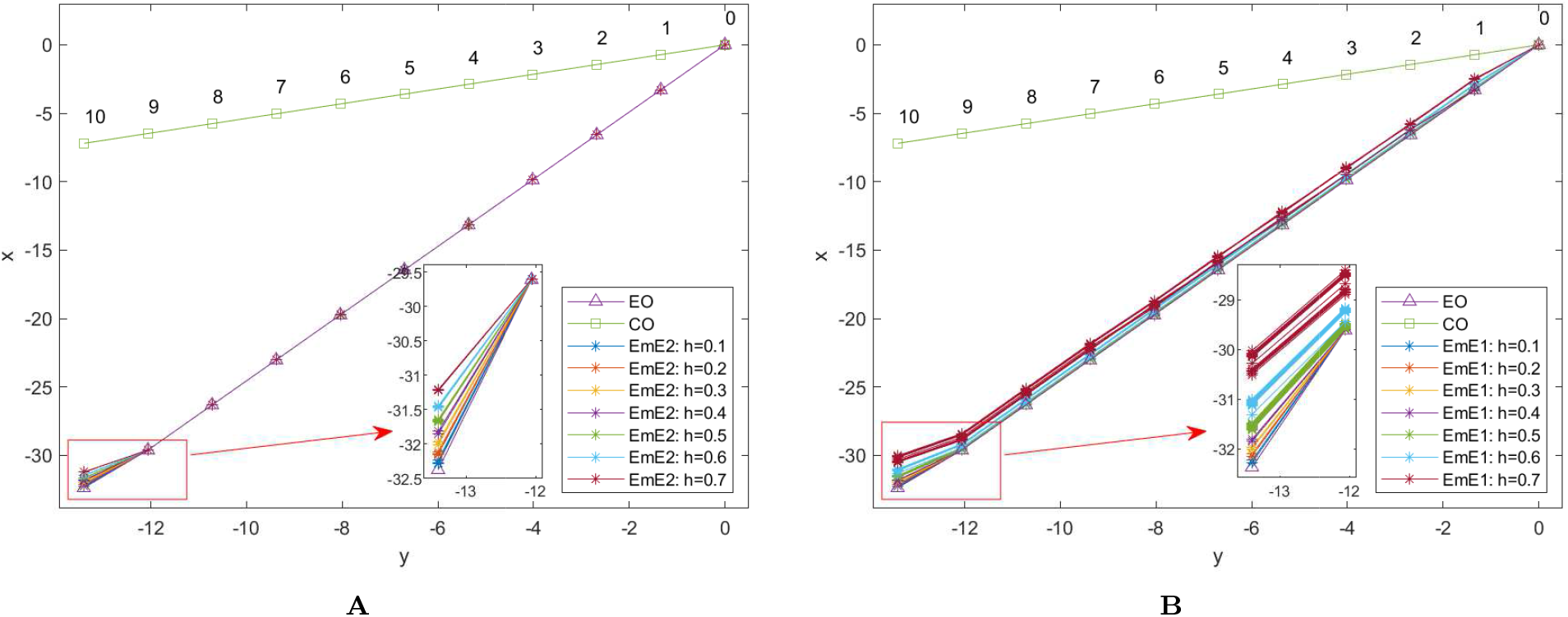
Formation for empathy flocks with empathy degree *h* = 0.1, …, 0.7 and the same simulation setting as in Fig. 7. **A**: Formation at the EmE2 (equilibrium when birds show empathy to both front and back neighbors) **B**: Formation at the EmE1 (equilibrium when birds only show empathy to front neighbor).

**S1 Table.**
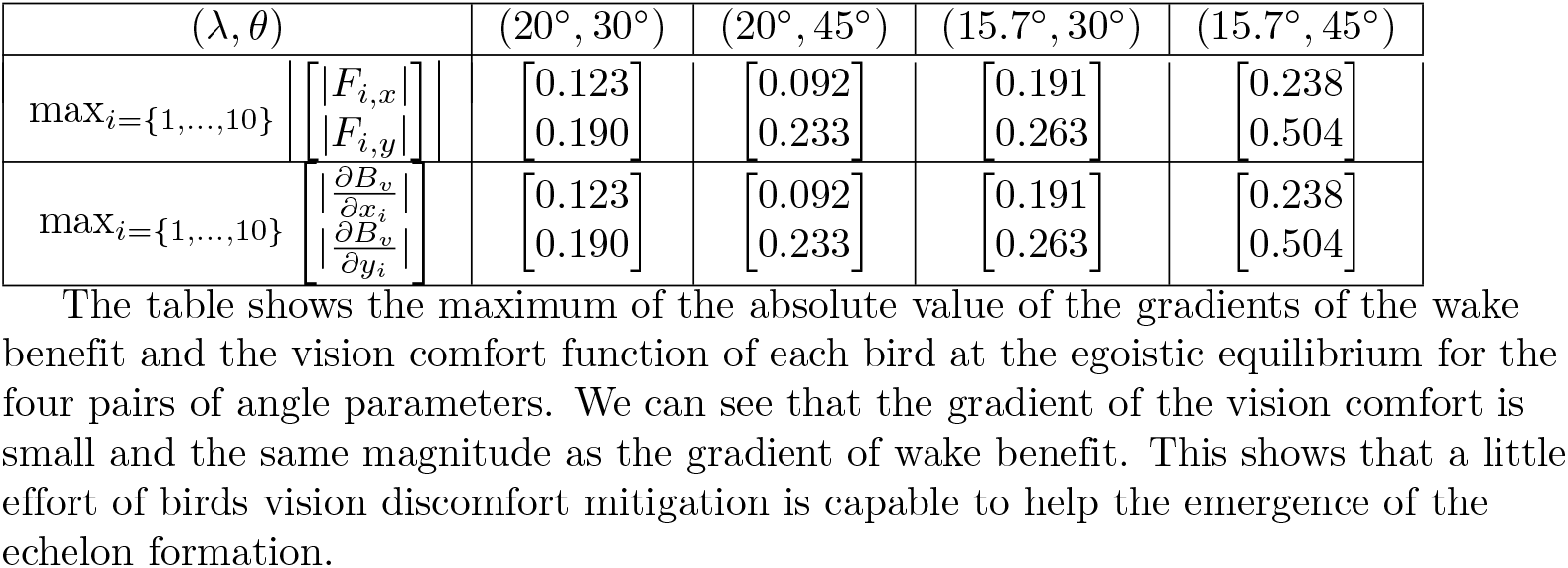

## Acknowledgments

The authors appreciate very much for the introducing and explaining of the aerodynamic wake model by our colleagues Ignace Ransquin and Philippe Chatelain at the Institute of Mechanics, Materials and Civil Engineering of UCLouvain. Computational resources have been partly provided by the supercomputing facilities of the Université catholique de Louvain (CISM/UCL) and the Consortium des Équipements de Calcul Intensif en Fédération Wallonie Bruxelles (CÉCI) funded by the Fond de la Recherche Scientifique de Belgique (F.R.S.-FNRS) under convention 2.5020.11 and by the Walloon Region.

